# Hydrogels Containing Gradients in Vascular Density Reveal Dose-Dependent Role of Angiocrine Cues on Stem Cell Behavior

**DOI:** 10.1101/2021.02.12.431015

**Authors:** Mai T. Ngo, Victoria R. Barnhouse, Aidan E. Gilchrist, Christine J. Hunter, Joy N. Hensold, Brendan A.C. Harley

**Affiliations:** Dept. Chemical and Biomolecular Engineering, University of Illinois at Urbana-Champaign, Urbana, IL 61801; Dept. Bioengineering, University of Illinois at Urbana-Champaign, Urbana, IL 61801; Dept. Materials Science and Engineering, University of Illinois at Urbana-Champaign, Urbana, IL 61801; Institute for Genomic Biology, University of Illinois at Urbana-Champaign, Urbana, IL 61801; Cancer Center at Illinois, University of Illinois at Urbana-Champaign, Urbana, IL 61801

**Keywords:** stem cells, hydrogels, vascularization, microfluidics, gradients, patterning

## Abstract

Biomaterials that replicate patterns of microenvironmental signals from the stem cell niche offer the potential to refine platforms to regulate stem cell behavior. While significant emphasis has been placed on understanding the effects of biophysical and biochemical cues on stem cell fate, vascular-derived or angiocrine cues offer an important alternative signaling axis for biomaterial-based stem cell platforms. Elucidating dose-dependent relationships between angiocrine cues and stem cell fate are largely intractable in animal models and two-dimensional cell culture. In this study, we leverage microfluidic mixing devices to generate three-dimensional hydrogels containing lateral gradients in vascular density alongside murine hematopoietic stem cells (HSCs). Regional differences in vascular density can be generated via embossed gradients in cell, matrix, or growth factor density. HSCs co-cultured alongside vascular gradients reveal spatial patterns of HSC phenotype in response to angiocrine signals. Notably, decreased Akt signaling in high vessel density regions led to increased expansion of lineage-positive hematopoietic cells. This approach offers a combinatorial tool to rapidly screen a continuum of microenvironments with varying vascular, biophysical, and biochemical cues to reveal the influence of local angiocrine signals on HSC fate.

## 1. Introduction

Regulation of stem cell behavior is essential for maintaining normal tissue function and repair, but it can also contribute to the onset and progression of disease.^[1]^ For example, balancing stem cell expansion while preventing depletion of the stem cell population is critical for maintaining a population that can facilitate tissue repair or regeneration when injury occurs.^[2]^ In some cancers, a subpopulation of cancer stem cells has been attributed with tumorigenicity and recurrence after initial treatment.^[3]^ Stem cells are influenced by their local niche, which includes the biophysical, biochemical, and cellular properties of the surrounding microenvironment.^[4]^ Thoroughly understanding how microenvironmental properties instruct stem cell behavior and dysregulation remains a central challenge for developing *ex vivo* culture platforms and scalable stem cell therapies.

The vascular niche is the local tissue microenvironment adjacent to vascular structures in tissues. Here, endothelial cells and perivascular stromal cells provide vascular-derived or angiocrine signals believed to play a central role in directing stem cell behavior.^[5]^ Hematopoietic stem cells (HSCs), for example, have been shown to reside in perivascular regions in which vascular cells provide signals that direct the balance between quiescence, self-renewal, and differentiation.^[5b, 6]^

Being able to correlate vascular architecture and density to stem cell outcomes provides enormous opportunities for engineering patterns of angiocrine-inspired signals into ex vivo platforms, but deriving such design principles is challenging. Traditional two-dimensional cultures lack the ability to recapitulate vascular architecture and the biophysical aspects of the vascular niche, which undergoes extensive remodeling during vasculogenic or angiogenic events.^[7]^ Dynamic remodeling of the microenvironment has been shown to profoundly impact stem cell behavior by affecting the balance between paracrine and autocrine signaling.^[8]^ While animal models are able to recapitulate the full complexity of the vascular niche, characterizing local vascular architecture and connecting such metrics with regional stem cell phenotype is challenging. Three-dimensional biomaterial platforms offer a robust approach for creating benchtop models of the vascular niche that recapitulate vascular development and architecture within a dynamically remodeling microenvironment.^[9]^

Complex patterns of biophysical, biochemical, and angiocrine cues within vascular niches suggest a large potential design space for exploring signal-response relationships between engineered niches and stem cell phenotype.^[10]^ The biomaterials community has recently developed a wide range of tools to generate combinatorial platforms that leverage photopatterning,^[11]^ microarrays,^[12]^ and liquid handling protocols.^[13]^ Our lab has previously developed microfluidic mixing devices that allow us to generate collagen and gelatin-based hydrogels containing lateral gradients of cell, matrix, and biomolecular signals.^[14]^ Local analysis of cell response from defined regions within these gradient hydrogels post-culture allows for quick mapping of cell phenotype in response to local changes in microenvironmental signals. Separately, we have developed technologies to incorporate endothelial and stromal cells into monolithic hydrogels to mimic aspects of the perivascular niche.^[15]^ However, the combination of these technologies (gradient hydrogels; vascular co-culture) has not yet been leveraged to generate combinatorial models of the perivascular microenvironment.

In this study, we report the successful development of a microfluidic device to generate hydrogels containing lateral gradients in vascular density as a means to identify angiocrine signals underlying hematopoietic stem cell culture ex vivo. We first identify multiple routes (cell density gradients; matrix density gradients; gradients of matrix-bound VEGF) to generate hydrogels containing vascular gradients. We then report the role of increasing vascular density in the balance of activated vascular cells and secretion of angiocrine factors that can influence the culture of primary murine HSCs, which results in differential capacity for large-scale expansion of lineage-committed hematopoietic cells versus local maintenance of a stem cell population as a function of vascular complexity.

## 2. Results

### 2.1. Microfluidic Mixing Devices Can Generate Hydrogels Containing Overlapping Gradients

We generated two classes of patterned hydrogel gradients: 1) opposing linear gradients of two factors; and 2) a linear gradient of one factor embossed on a constant density of a second factor. Gradients were visualized via blue and red fluorescent beads mixed into the prepolymer solution prior to loading into the microfluidic device (**Figure 1A**). First, quantitative analysis of blue and red channel fluorescent intensity across the resulting hydrogels revealed the ability to generate hydrogels with opposing fluorescent gradients (**Figure 1B**). We subsequently generated hydrogels containing a single lateral gradient (red) embossed across a hydrogel containing a constant density of a second factor (blue) (**Figure 1C**). Quantitative analysis of local fluorescent intensity confirmed a constant fluorescent intensity in the blue channel, but a lateral gradient in crimson bead fluorescent intensity. Taken together, this data demonstrates that the microfluidic mixing device can generate overlapping gradients of multiple stimuli based on the geometric mixing of precursor inlet suspensions.

**Figure 1.**
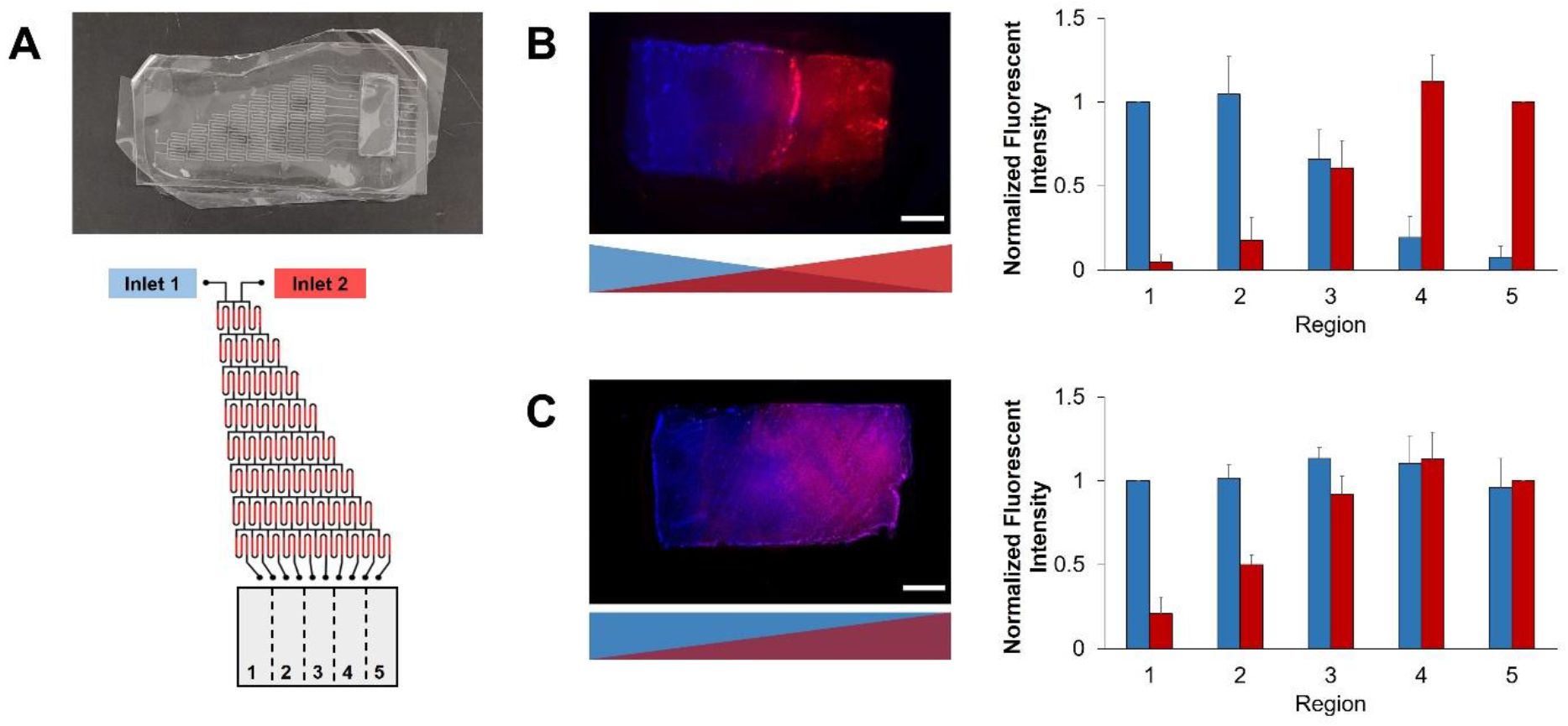
**(A)** Schematic of the microfluidic mixing device. **(B)** and **(C)** Fluorescent beads are used to demonstrate that overlapping patterns of microenvironment cues can be patterned using the microfluidic mixer. N = 5 devices

### 2.2 Gradient Vascularization Generated by Differential Cell Density

We subsequently generated hydrogels containing gradients in initial vascular cell density and evaluated the influence of cell density on resultant vessel network formation over seven days. Hydrogels were generated by mixing two pre-polymer solutions, one containing human umbilical vein endothelial cells (HUVECs) and mesenchymal stromal cells (MSCs) while the second pre-polymer solution contained no cells. (**Figure 2A**). We generated variants with different ratios of HUVECs and MSCs (2:1, 4:1, 5:1 HUVEC:MSC) while keeping the overall density of HUVECs constant (2 × 10^6^ HUVECs/mL). After seven days of culture, immunofluorescent staining for CD31 revealed varying extents of vascular network formation across the hydrogel. Extensive vascular network formation was observed in regions of high initial cell density, with vascular network density gradually diminishing in regions with reduced cell density (**Figure 2B**). Analysis of vessel network complexity using metrics of branches/junctions, average branch length, and total network length further confirmed the presence of a gradient of vascular network formation across the hydrogels (**Figure 2C**, **S1**). Interestingly, the resolution of the gradient, assessed here by the presence of statistically significant differences in metrics between hydrogel regions, was dependent upon HUVEC:MSC ratio. While there were no differences in vascular network metrics between regions at a 5:1 HUVEC:MSC ratio, statistically significant differences were observed for hydrogels containing 2:1 and 4:1 HUVEC:MSC ratios. Specifically, total network length and number of branches and junctions differed between regions in 2:1 HUVEC:MSC hydrogels, while all metrics differed between regions in 4:1 HUVEC:MSC hydrogels. Overall, these results demonstrate that generating initial gradients in the local density of endothelial and stromal cells provides an avenue to locally control formation of regionally distinct vascular features across a single hydrogel.

**Figure 2.**
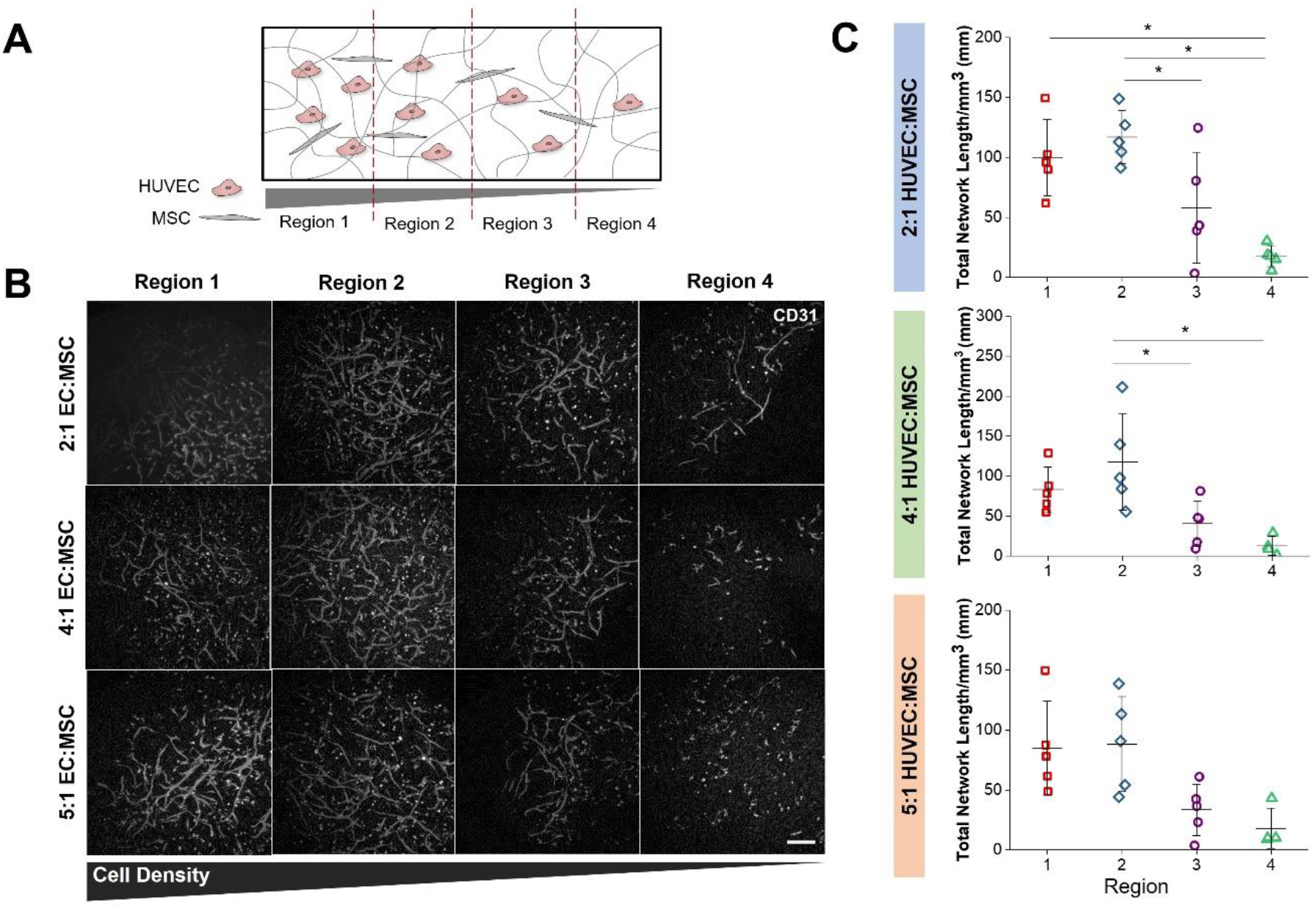
**(A)** A gradient of vascular cell density is generated across GelMA hydrogels. **(B)** Visual inspection of the resulting vascular networks after seven days of culture reveals differences in regional vascularization with various HUVEC:MSC ratios. Scale bar = 200 μm. **(C)** Quantification of network length per region reveals that only 2:1 and 4:1 HUVEC:MSC ratios yields statistically significant differences in regional vascularization across the hydrogel. *p<0.05, N = 5 devices

### 2.3 Gradient Vascularization Generated by Manipulating the Hydrogel Microenvironment

Because the extent of vascularization is tightly correlated to the biophysical and biochemical properties of the surrounding microenvironment,^[16]^ we hypothesized that hydrogels containing gradients in vascular network formation could be generated by manipulating the properties of the hydrogel itself. We first generated hydrogels containing a constant density of vascular cells but a lateral gradient of matrix density (opposing inlet suspension of 5 wt% vs. 7 wt% GelMA solutions; **Figure 3A, 3B**). All metrics of vascular complexity demonstrated a decrease in vessel formation with increasing matrix density (**Figure 3C**, **S2**). In particular, statistical significance was observed between Regions 1 and 2 (low matrix density) compared to Region 4 (high matrix density). We subsequently generated hydrogels containing a constant density of cells but a lateral gradient of covalently-bound VEGF (**Figure 3D**). Acrylate-PEG-VEGF was synthesized to tether VEGF to the hydrogel backbone and showed at least 75% retention in GelMA over 48 hours (**Figure S3**). Vessel network complexity trended upwards with increasing VEGF tethering, with statistically significant differences observed between Regions 1 and 4 (**Figure 3E, 3F, S3**).

**Figure 3.**
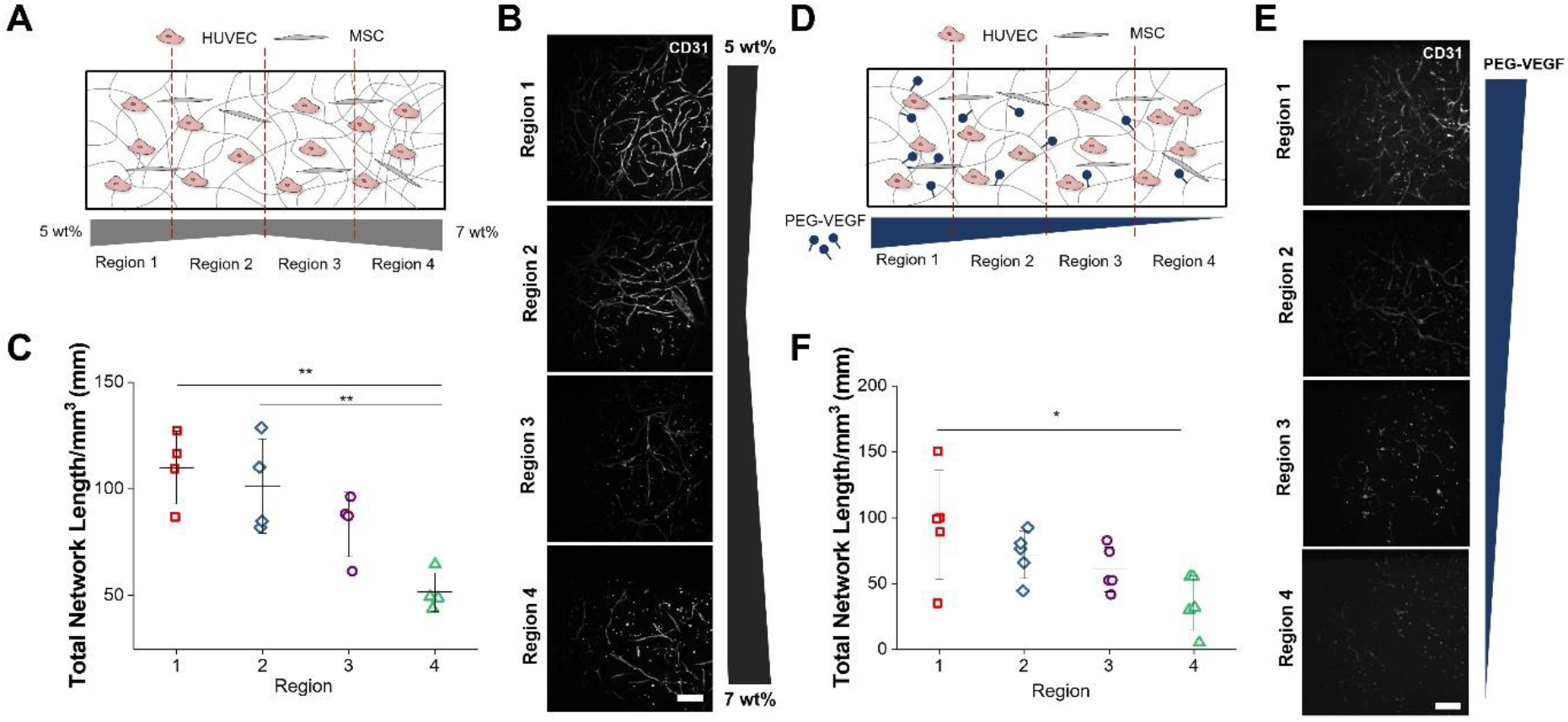
**(A)** Hydrogels containing a gradient in matrix density were generated by mixing 5 and 7 wt% hydrogel solutions with equivalent vascular cell densities. **(B)** Qualitative and **(C)** quantitative analysis reveals the presence of a gradient in the extent of vascularization as a function of varying matrix density. **p<0.01, N = 4 devices **(D)** Hydrogels were generated with a gradient in covalently-bound VEGF and a constant density of vascular cells. **(E)** Qualitative and **(F)** quantitative analysis reveals the presence of a gradient in the extent of vascularization as a function of VEGF functionalization. *p<0.05, N = 5 devices

### 2.4 Angiocrine Signals Support Expansion of Differentiated Hematopoietic Population While Maintaining a Pool of Undifferentiated Stem Cells

We next examined the potential application of hydrogels containing lateral vascular gradients to locally regulate cell phenotype. We fabricated hydrogels containing a lateral vessel gradient (4:1 HUVEC:MSC) but a constant distribution of HSCs (1 × 10^5^/mL) (**Figure 4A**). After seven days, hydrogels were fixed and stained with CD45 and CD31 to visualize hematopoietic cells and vasculature respectively (**Figure 4B**). Clusters of round, CD45+ hematopoietic cells were observed in proximity to vasculature in regions of high vascular density. Additional hydrogels were divided into four sections, with encapsulated cells extracted via collagenase degradation and subsequently assessed via flow cytometry for the presence of surface markers indicating stem-like or differentiated status (**Figure 5A**). While there were no statistically significant differences in the number of stem cells as a function of vascular cell density (**Figure 5B**), the number of differentiated, lineage-positive cells increased with increasing vascular density (**Figure 5C** and **S4**). Statistically significant differences were observed between Regions 1 (High vascular cell density) and 3, and between all regions compared to Region 4 (Low vascular cell density). We additionally analyzed the fraction of quiescent HSCs per region, with quiescence defined by the G0 phase of the cell cycle (Ki-67^−^, DAPI^≤2N^). While not statistically significant, the number of quiescent HSCs trended upwards with the increasing presence of vascular cells (**Figure 5D**).

**Figure 4.**
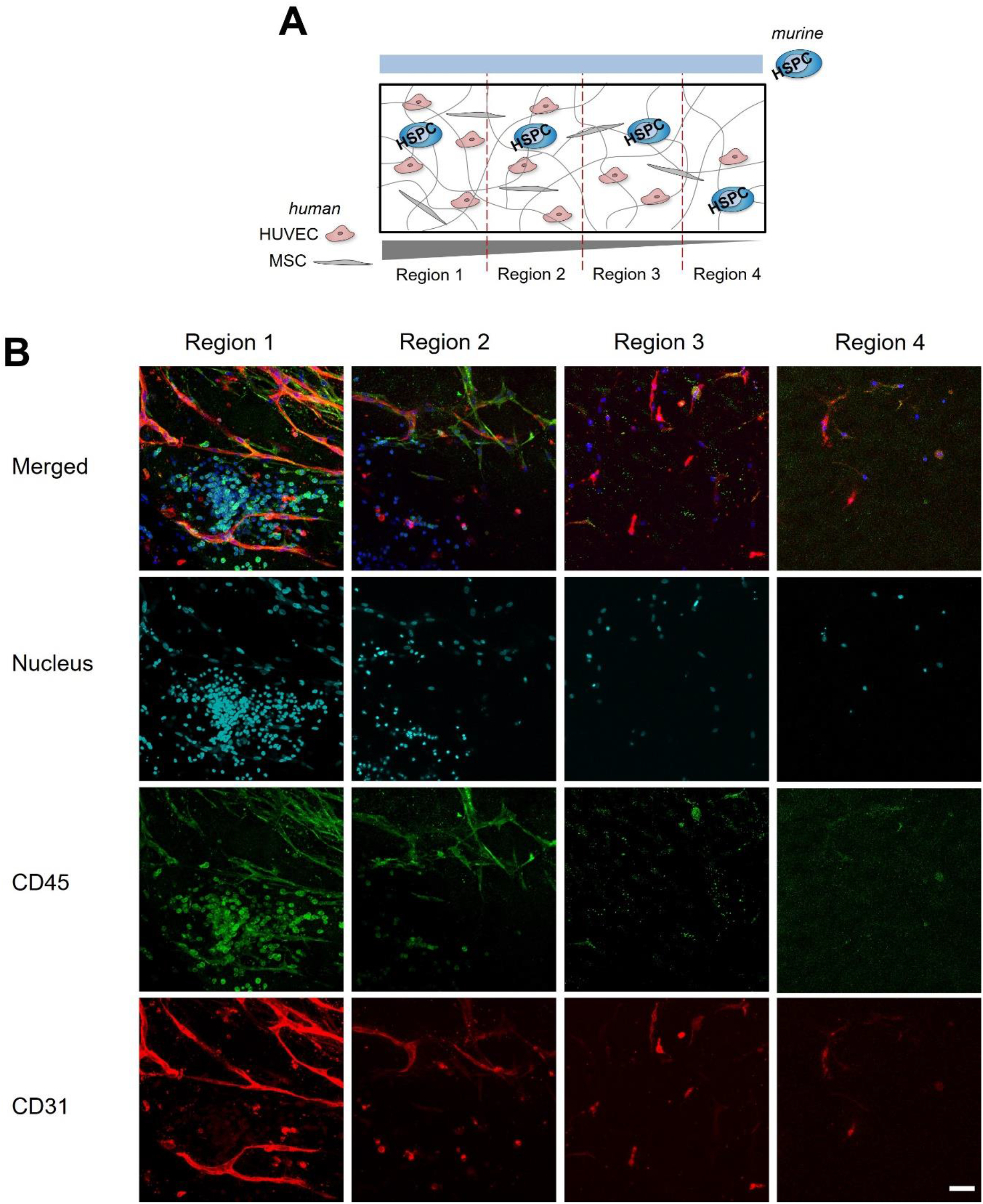
**(A)** Hydrogels were established with a gradient of human vascular cells and a constant presence of murine-derived hematopoietic stem and progenitor cells (HSPCs). **(B)** Representative staining for CD31 (vasculature) and CD45 (hematopoietic cells) across the hydrogel regions. Scale bar: 50 μm

**Figure 5.**
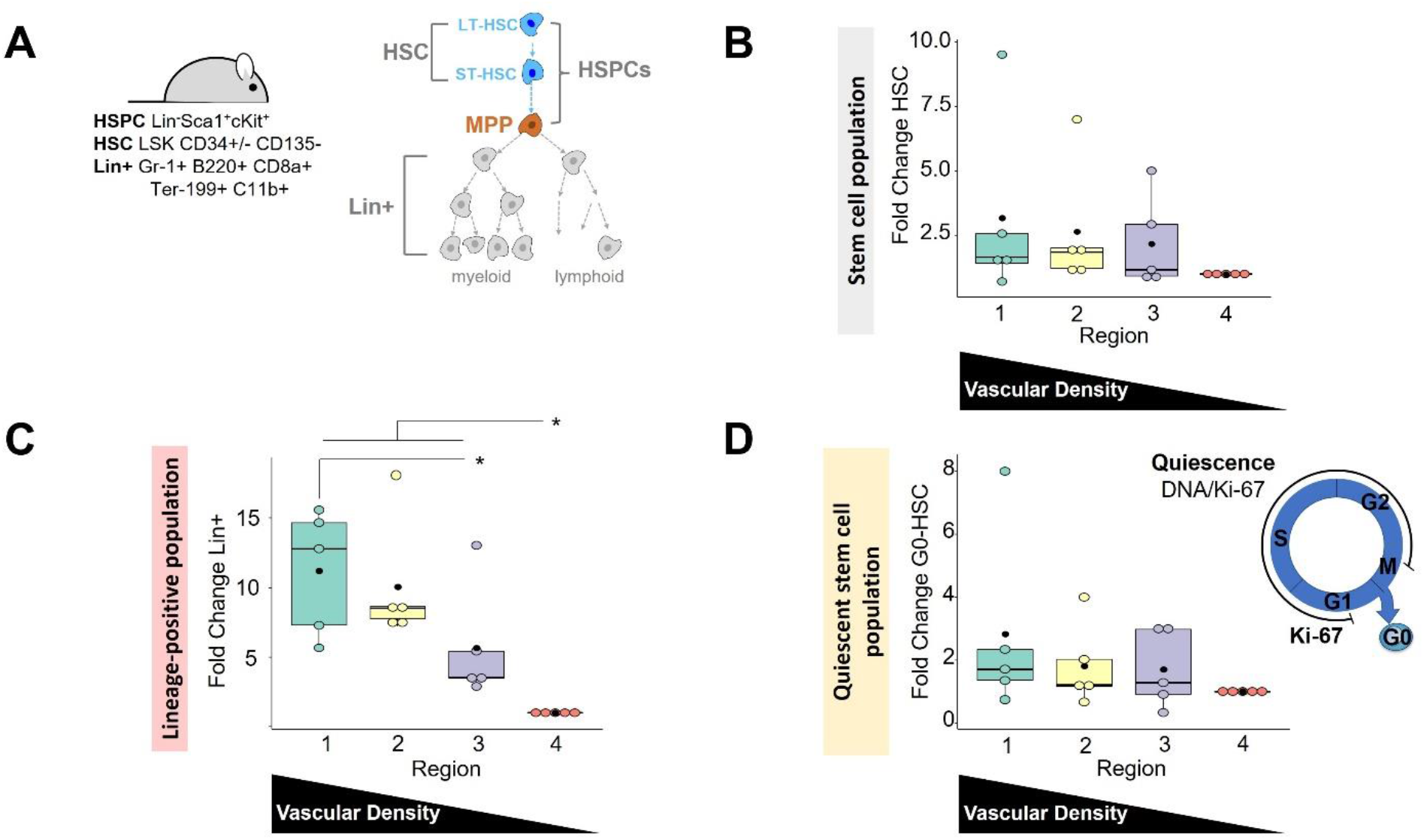
**(A)** Surface marker staining and flow cytometry was used to identify hematopoietic stem cells (HSCs) and lineage-positive hematopoietic cells in the different regions of the hydrogel. **(B)** The number of HSCs did not vary between regions. **(C)** The number of lineage-positive hematopoietic cells significantly increased with increasing vascular density. **(D)** The fraction of HSCs that were quiescent did not vary between hydrogel regions. In all graphs, the number of cells of interest per region are normalized to the number of cells of interest in Region 4. *p<0.05, N = 5 devices

### 2.5 Increasing Vascular Cell Density Correlates with Decreasing Akt Activation and Increased ANG2 Secretion

To examine potential mechanisms by which vascular density regulates hematopoietic stem cell fate in a dose-dependent manner, we examined the activation of MAPK and Akt signaling pathways in HUVECs and MSCs using a series of monolithic (non-gradient) hydrogels fabricated using a range of cell densities meant to simulate the transitions across the spatially-graded hydrogels (5 × 10^5^ HUVEC/mL, 1 × 10^6^ HUVEC/mL, 2 × 10^6^ HUVEC/mL; 4:1 HUVEC:MSC). Phosphorylated to total (p/t)AKT decreased signficantly with increasing vascular density between 5 × 10^5^ HUVEC/mL and 2 × 10^6^ HUVEC/mL groups (**Figure 6A**, **6B**, **S5**). We subsequently used a cytokine array to compare the secretome between hydrogels as a funciton of vascular density (**Figure 6C**). ANG2 secretion demonstrated a statistically significant increase at the highest density (2 × 10^6^ HUVEC/mL) compared to 5 × 10^5^ HUVEC/mL and 1 × 10^6^ HUVEC/mL densities. IGFBP2 and TSP1 secretion also trended upwards with increasing vascular density. Finally, immunofluorescent staining revealed Jagged1 expression by vascular networks was not affected by local vascular density (**Figure 6D**).

**Figure 6.**
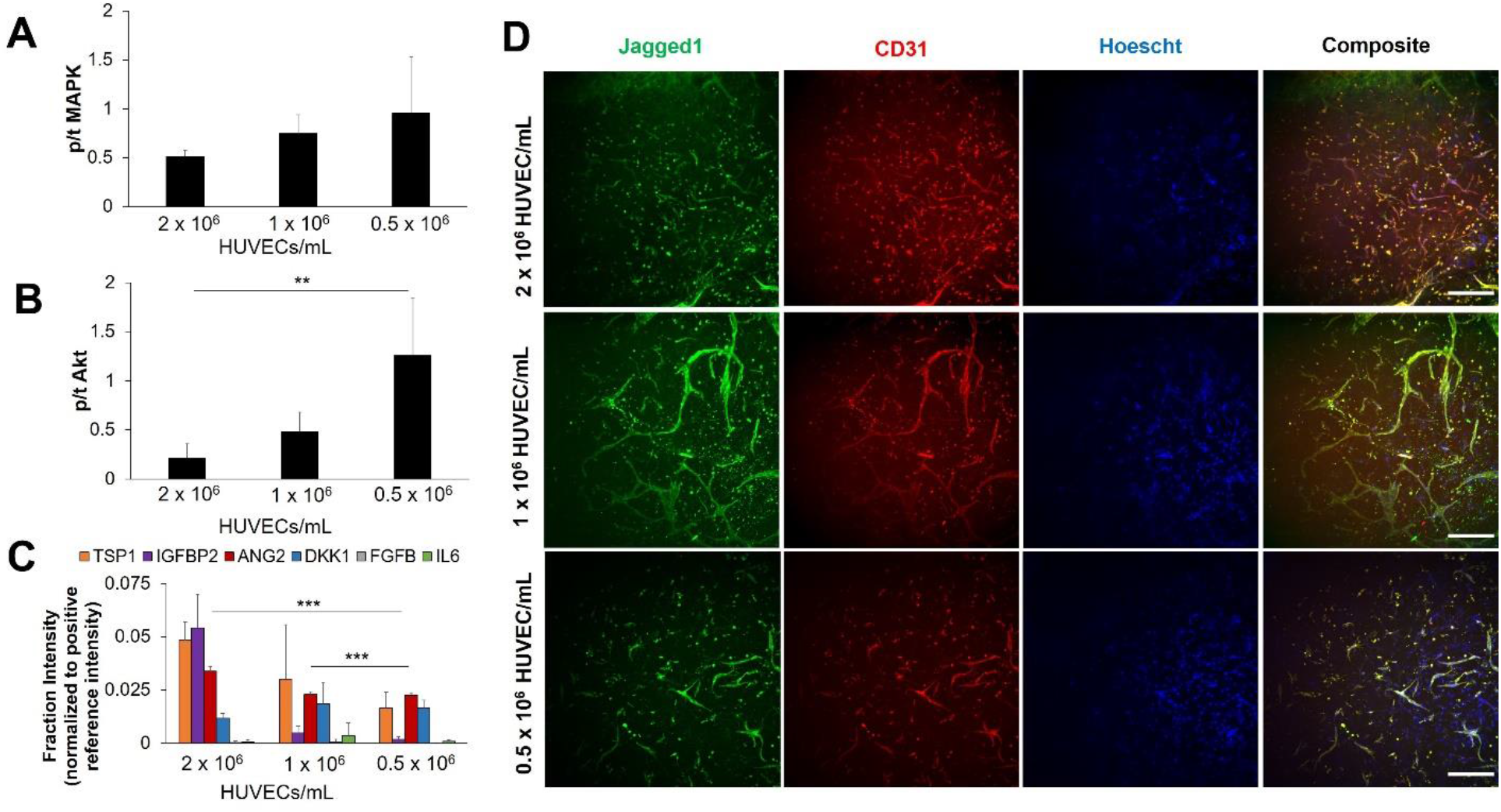
**(A)** and **(B)** Quantification of MAPK and Akt pathway activation measured using Western blot. **p<0.01, N = 5 hydrogels **(C)** A cytokine array was used to compare the abundance of secreted proteins between hydrogels with different vascular densities. ***p<0.001, N = 3 conditioned media samples **(D)** Vascular networks express Jagged1 across all cell densities. Scale bar = 200 μm

## 3. Discussion

Stem cell behavior is controlled by the collective influence of biophysical, biochemical and cellular cues presented by the niche microenvironment.^[17]^ Identifying the individual and synergistic effects of the various microenvironmental properties on stem cell phenotype is important for developing therapies for regenerative medicine and understanding the mechanisms that initiate diseases such as cancer. However, the vast complexity of the niche microenvironment creates the need for high-throughput or combinatorial technologies to efficiently screen through various combinations or dosages of microenvironmental properties and characterize the resultant stem cell phenotype. While various platforms have been developed over the past several years for the purpose of high-throughput screening of stem cell behavior,^[18]^ the inclusion of vascular cues has not yet been explored. The vascular stem cell niche has been shown to influence neural and hematopoeitic stem cell behavior,^[19]^ and additionally contributes to tumor maligancy by regulating cell dormancy,^[20]^ migration,^[21]^ and therapeutic response.^[22]^ Within the field of regenerative medicine, there is enormous interest in promoting vascularization alongside stem cell activity in therapuetic platforms because of the need to supply nutrients and oxygen to the regenerating or repaired tissue.^[23]^ However, vascular cells provide additional cues through angiocrine signalling that influence stem cell behavior.^[5c]^ Furthermore, active angiogenic or vasculogenic processes leads to extracellular matrix remodeling, and dynamic biophysical and biochemical changes to the microenvironment have been shown to influence stem cell fate.^[8a, 24]^ Thus, an understanding of how the recipricocity between the niche microenvironment and vascular processes controls stem cell fate is critical for accurately defining how the vascular niche contributes to disease progression and stem cell therapies.

Overall, our results show that hydrogels containing gradients in vascular density can generate spatially-graded angiocrine environments sufficient to locally regulate stem cell phenotype. We used microfluidic mixing devices to generate hydrogels containing gradients in vascular density. Microfluidic mixing devices are easy to fabricate using standard soft lithography approaches, enable rapid generation of graded biomaterial environments, and allow for the exploration of continuous ranges of microenvironmental variables.^[25]^ We and other groups have demonstrated that vascularization of three-dimensional biomaterials is dependent upon cell density,^[26]^ matrix density or stiffness,^[27]^ and growth factor presentation;^[28]^ thus, we explored the ability to continuously alter vascular density across single hydrogels by generating gradients of these various parameters. Our results revealed the ability to generate distinct regional differences in vascular density across single hydrogels via gradients in vascular cell density, matrix density, and VEGF presentation. We additionally demonstrated that variation in the extent of vascularization was dependent on the ratio of endothelial to stromal cells, likely reflecting the role of stromal cells in tuning vascularization by secreting angiogenic growth factors or facilitating matrix remodeling.^[29]^ The facile nature of our approach allows future exploration of growth factor gradients of varying steepness to achieve significant regional differences in vascularization. The ability to immobilize growth factors in a spatial manner also enables investigation into the reciprocal effects of the biomolecular microenvironment on vascular niche development and stem cell behavior. With the vascularization of engineered tissues remaining a signficant challenge in regenerative medicine,^[30]^ our results also suggest that gradient hydrogels can be used as a tool to optimize biomaterial parameters to achieve appropriate amounts of vascularization.

To demonstrate how our platform can be used to evaluate the role of angiocrine cues in influencing stem cell behavior in a dose-dependent manner, we examined hematopoeitic stem cell (HSCs) fate as a function of local vascular density. Tens of thousands of HSC transplantations are performed annually and are used to treat diseases such as leukemia, lymphoma, and autoimmune disorders.^[31]^ However, graft failure is a significant clinical challenge.^[32]^ While expanding the number of transplanted cells is a promising solution for improving successful engraftment, the ability to expand functional HSCs *ex vivo* has proven to be difficult.^[33]^ Biomaterial platforms that are able to direct HSC expansion while also retaining a fraction of quiescent long-term repopulating HSCs remains a clinical need,^[34]^ and despite the well-established role of the vascular niche in regulating HSC behavior,^[6a, 15a, 19a, 35]^ the use of angiocrine-inspired cues in biomaterials for HSC culture has yet to be fully explored. Here, we show increasing local presence of HUVECs and MSCs dramatically increases the expansion of differentiated hematopoieitic cells but not at the expense of maintaining a subpopulation of HSCs. This general result is consistent with *Braham et al.*, who demonstrated that co-culturing HSCs with mesenchymal stromal and endothelial progenitor cells in alginate or Matrigel promoted hematopoeitic proliferation;^[36]^ however, our platform provides the first evidence that vascular density plays a dose-dependent role in modulating HSC behavior. Infusion of endothelial progenitor cells has been shown to improve hematopoieitic stem cell reconstitution, partly due to the re-establishment of a vascular niche environment post-radiation.^[37]^ Our results suggest the opportunity to exploit *ex vivo* culture platforms that mimic angiocrine signals for expansion of differentiated cells important for short-term repopulation of the hematopoeitic pool post-transplantation.^[38]^ Most importantly, these cultures suggest a wide range of avenues to refine the role of angiocrine cues on HSC lienage specification vs. expansion.

Recent work from *Kobayashi et al*. demonstrated that Akt-activated endothelial cells promote HSC and progenitor expansion while MAPK activation or co-activation with Akt favors the maintenance of the stem and progenitor pool and the expansion of differentiated cells.^[6b]^ Based on our findings of angiocrine environments locally altering hematopoietic expansion, we took advantage of the opportunity to use defined hydrogel cultures to compare MAPK and Akt activation as well as secretomic output as a function of vascular density. Excitingly, our multi-dimensional hydrogel platforms mimics the phenoptype of Akt activation decreasing with increasing vascular density, which may lead to an angiocrine output predominantly influenced by MAPK signaling to support the expansion of a lineage-positive hematopoieitic population. Observed increases in ANG2 secretion in hydrogels containing increased vascular density may be a result of dominant MAPK signaling, with literature demonstrating that ANG2 supports hematopoeitic cell proliferation but not long-term reconstitution.^[39]^ ANG2 also antagonizes ANG1-induced phosphorylation of Akt, further decreasing Akt activation.^[39b]^ The retention of a HSC subpopulation irrespective of vascular density may be supported by IGFBP2 secretion along with the expression of the Notch ligand Jagged1 by the vascular networks, which collectively have been shown to regulate HSC survival, maintenance, and cycling.^[40]^ Future exploration that stems from this work will focus on the establishment of hydrogels containing gradients in covalently-tethered angiopoietins, IGFBP2, and Notch ligands individually and in synergy to further isolate the roles of these biomolecules in modulating *ex vivo* HSC fate. Another interesting avenue involves the generation of gradient vascular hydrogels using Akt-activated, *E4ORF1*-transduced endothelial cells to potentially shift the HSC response towards stem and progenitor cell expansion.^[6b]^ Finally, while the linear gradients demonstrated here have advantages for showing the feasibility of vascular gradations influencing hematopoietic cell activity, the complexity of the bone marrow microenvironments suggests the need for more complex patterns of vascular density. Ongoing efforts in our laboratory are examining modifications to the microfluidic mixing device to alter the shape and steepness of the resulting gradients in order to improve the study of regional stem cell behavior across the platform as a function of the local microenvironment.

## 4. Conclusion

In this work, we describe the generation of hydrogels containing gradients in vascular density by combining microfluidic technologies with design principles governing angiogenic processes in biomaterials. Application of this platform to understand the dose-dependent role of angiocrine signaling on HSC fate reveals that increasing vascular density differentially modulates the expansion of differentiated hematopoietic cells without exhaustion of a stem and progenitor cell subpopulation, with these stem cell fate decisions potentially driven by MAPK over Akt activation and enhanced ANG2 secretion by engineered vessel networks. We anticipate that this platform can be used to explore or optimize the presentation of angiocrine cues for regenerative therapies with other stem cells (e.g. neural stem cells) as well as advance our understanding of the role of angiocrine signals in disease progression.

## 5. Experimental Section/Methods

### Microfluidic Mixer Fabrication

The microfluidic mixer device was fabricated as previously described.^[41]^ Briefly, standard photolithography approaches were used to generate a master mold with a negative relief of the device on a silicon wafer (University Wafer, South Boston MA). The device contained eight rows of 200 μm x 100 μm channels, with each channel containing 50 μm tall, staggered herringbone features to facilitate chaotic advection and mixing within the channels. The channels empty into a rectangular well to form a hydrogel. To fabricate the master mold, a 100 μm thick layer of SU8-2050 photoresist was spin-coated on the wafer, baked, and then exposed to UV light after the application of a photomask containing the channel design. A second 50 μm thick layer was then spin-coated on the wafer, baked, and exposed to UV light after the application of a second photomask containing the herringbone features. The photoresist was then baked and developed using propylene glycol monomethyl ethyl acetate (PGMEA, Sigma Aldrich, St. Louis, MO). Pieces of silicon wafer were glued to the portion of the master mold containing the rectangular well to create a 1 mm thick well region. The resulting mold was silanized using (heptadecafluoro-1,1,2,2-tetrahydrodecyl)trimethoxysilane (Gelest, Morrisville, PA) to prevent polydimethylsiloxane (PDMS) from sticking to the master mold. PDMS monomer and crosslinker (Sylgard 184, Dow Chemical Company, Midland, MI) were mixed in a 5:1 ratio, poured onto the master mold, and crosslinked for 1 hour at 65 °C. After the PDMS was removed from the master mold, inlet ports were punched into the mixing device using a needle. On a separate piece of non-patterned PDMS, a rectangular punch was used to punch out a rectangle with the same dimensions as the well in the mixing device. The PDMS pieces along with a transparent polycarbonate membrane (MilliporeSigma, Burlington, MA) were cleaned with oxygen plasma and submerged in (3-Aminopropyl)triethoxysilane (APTMS, Sigma Aldrich, St. Louis, MO) and (3-Glycidyloxypropyl)triethoxysilane (GPTMS, Sigma Aldrich, St. Louis, MO) respectively. The final device was constructed by bonding the PDMS pieces with the membrane in between, such that the rectangular well on the patterned PDMS piece was aligned with the rectangular hole on the blank piece of PDMS (**Figure 1A**). The device was dried overnight at 65 °C before use.

### Vascular Cell Culture

Human umbilical vein endothelial cells (HUVECs) and human bone marrow-derived mesenchymal stromal cells (MSCs) were purchased from Lonza (Walkersville, MD). HUVECs were cultured using Endothelial Growth Medium 2 (EGM2, Lonza, Walkersville, MD) while MSCs were cultured using DMEM supplemented with penicillin/streptomycin and FBS (mesenchymal stromal cell qualified, Thermo Fisher Scientific, Waltham, MA). All media were further supplemented with plasmocin prophylactic (Invivogen, San Diego, CA) to prevent mycoplasma contamination. All cells were used before passage five and were cultured at 37 °C and 5% CO_2_.

### Primary HSC Isolation from Murine Bone

All work involving primary cell isolation was conducted under approved animal welfare protocols (Animal Protocol #20078, Institutional Animal Care and Use Committee, University of Illinois at Urbana-Champaign). Bone marrow cells were isolated from the crushed tibia and femur of C57BL/6 female mice, age 4 – 8 weeks (The Jackson Laboratory, Bar Harbor, ME). A suspension of whole bone marrow cells was lysed with ACK buffer on ice for a maximum of 5 minutes to eliminate red blood cells. Cells were washed in PBS + 5% FBS at 300 rcf for 10 minutes. Initial hematopoietic lineage negative enrichment was performed with EasySep™ Mouse Hematopoietic Progenitor Cell Isolation Kit (#19856, Stemcell Technologies, Vancouver, Canada). The enriched lineage-negative (Lin-) population were then stained for further purification via fluorescence-activated cell sorting (FACS). A BD FACS Aria II flow cytometer was used to collect the LSK population: Lin^−^ Sca^−^1^+^ c-kit^+^, with propidium iodide (#BMS500PI, Thermo Fisher Scientific, Waltham, MA) for dead cell exclusion. LSK antibodies were supplied by eBioscience (San Diego, CA), and are as follows: APC-efluor780-conjugated c-kit (1:160, #47-1172-81), PE-conjugated Sca-1 (0.3:100, #12-5981-83), and Lin: FITC-conjugated CD5, B220, CD8a, CD11b (1:100, #11-0051-82, #11-0452-82, #11-0081-82, #11-0112-82), Gr-1 (1:400, #11-5931-82), and Ter-119 (1:200, #11-5921-82).^[42]^

### GelMA Synthesis

Methacrylamide-functionalized gelatin (GelMA) was synthesized by dissolving 2 g of gelatin (porcine type A, 300 bloom, Sigma Aldrich, St. Louis, MO) in PBS at 60 °C. 250 μL of methacrylic anhydride (Sigma Aldrich, St. Louis, MO) was added dropwise and the reaction was allowed to proceed for one hour with stirring at 400 RPM. The reaction mixture was quenched with PBS, and the resulting solution was transferred to dialysis membranes (12-14 kDa cutoff, Thermo Fisher Scientific, Waltham, MA) and dialyzed against deionized water for seven days with daily solvent exchanges. The reaction mixture was then frozen and lyophilized, and the degree of functionalization was assessed to be ~60% via ^1^H NMR.

### PEG-VEGF Synthesis and Secretion

Acrylate-PEG-VEGF was synthesized as previously described.^[26]^ Briefly, 25 μg of recombinant VEGF_165_ (Peprotech, Rocky Hill, NJ) was reacted with acrylate-PEG-succininmidyl carboxymethyl ester (MW 3500, Sigma Aldrich, St. Louis, MO) in a 200:1 molar acrylate-PEG-NHS:VEGF ratio in PBS (pH = 8.0) for two hours at 4 °C on a rocker platform. The reaction was quenched with PBS (pH = 8.0) and dialyzed against PBS (pH = 7.4) overnight. 0.1% BSA was added to the reaction mixture before aliquoting and storage at −20 °C. Successful conjugation was determined using Western blot (**Figure S3**), in which SDS-PAGE was performed on VEGF and PEG-VEGF samples using a 4-20% Mini-PROTEAN TGX precast protein gel (Biorad, Hercules, CA) under reducing conditions. Goat anti-human VEGF_165_ and HRP-conjugated anti-goat antibodies (both R&D Systems, Minneapolis, MN) were used to probe for protein, and the SuperSignal West Pico Chemiluminescent Substrate kit (Thermo Fisher Scientific, Waltham, MA) was used for detection by an ImageQuant LAS 4000 (GE Healthcare Bio-Sciences, Pittsburgh, PA). Secretion kinetics for PEG-VEGF was determined by fabricating 5 wt% GelMA hydrogels containing either 2 ng/mL soluble VEGF or PEG-VEGF. Hydrogels were incubated in 500 μL PBS with 1% FBS, and PBS was collected and replaced at 12, 24, and 48 hours. The concentration of VEGF in the PBS at various time points was determined using ELISA (R&D Systems, Minneapolis, MN) and normalized to the starting VEGF concentration.

### Fluorescent Gradient Hydrogel Fabrication

5 wt% GelMA was dissolved in PBS at 65 °C, and lithium acylphosphinate (0.1% w/v) was added to generate the pre-polymer solution. Red and blue fluorescent beads (1 μm diameter, Thermo Fisher Scientific, Waltham, MA) were added to hydrogel pre-polymer solutions in order to visualize distinct classes of gradient devices. Hydrogels containing a single gradient of red beads across a constant density of blue beads were visualized using blue beads added to one pre-polymer solution (Inlet 1 in **Figure 1A**) while red and blue beads were added to the opposing pre-polymer solution (Inlet 2 in **Figure 1A**). Alternatively, opposing gradients were generated using blue beads added to one pre-polymer solution (Inlet 1 in **Figure 1A**) while red beads were added to the other pre-polymer solution (Inlet 2 in **Figure 1A**). Each solution was withdrawn into 1 mL syringes using micro-syringe pumps (Harvard Apparatus, Holliston, MA). Tubing connected to each syringe was then inserted into the inlet ports of the mixing device, and pre-polymer solution was perfused through the device at 40 μl/min. The pre-polymer solution was polymerized by placing the device under UV light (λ = 365 nm, 7.8 mW/cm^2^) for 30 seconds. The hydrogel was then extracted from the device and imaged as described below.

### Vascular Gradient Hydrogel Fabrication

GelMA pre-polymer solution was generated as described previously. To generate hydrogels containing gradients in cell density, 2:1, 4:1, or 5:1 HUVEC:MSC (2 × 10^6^ HUVECs/mL) were suspended in one portion of pre-polymer solution, while a second pre-polymer solution did not contain any cells. To create hydrogels containing a gradient in matrix density, 2:1 HUVEC:MSC (2 × 10^6^ HUVECs/mL) were suspended in 5 wt% and 7 wt% GelMA solutions. To create hydrogels containing a gradient in covalently-tethered VEGF, 2:1 HUVEC:MSC (2 × 10^6^ HUVECs/mL) were suspended in 5 wt% GelMA solutions, with one solution containing 2 ng/mL acrylate-PEG-VEGF. Hydrogels were fabricated using the mixing device as described in the previous section. Extracted hydrogels with gradients in cell and matrix density were cultured in 24-well plates for seven days in EGM2, while hydrogels containing PEG-VEGF were cultured in EGM2 without soluble VEGF.

### Fabrication of Individual Hydrogels with Varying Vascular Densities

GelMA pre-polymer solution was generated as described above. HUVECs and MSCs were resuspended in 5 wt% pre-polymer solution at a ratio of 4:1 HUVEC:MSC with varying total cell densities of 2.5×10^6^ cells/mL, 1.25×10^6^ cells/mL and 0.625×10^6^ cells/mL. Hydrogels were polymerized with UV light (λ = 365 nm, 7.14 mW/cm^2^) in a 20 μL-capacity Teflon mold, then cultured in 1:1 EGM2 : SFEM for seven days. Hydrogels were collected for further analysis by Western blot and immunofluorescent staining. A subset of hydrogels (4 hydrogels per cell density) were cultured in serum and growth factor free endothelial basal media (EBM) for the final 24 hours of culture, after which the media was collected for analysis.

### HSC-Vascular Gradient Hydrogel Fabrication

HUVECs (2 × 10^6^ HUVECs/mL), MSCs (5 × 10^5^ MSCs/mL), and HSCs (1 × 10^5^ HSCs/mL) were suspended in one portion of pre-polymer solution, while only HSCs (1 × 10^5^ HSCs/mL) were suspended in a second portion of pre-polymer solution. Hydrogels containing vascular cell density gradients and a constant distribution of HSCs were cultured for seven days in 1:1 EGM2 : SFEM, with SFEM (Stemcell Technologies, Vancouver, Canada) containing 100 ng/mL SCF (Peprotech, Rocky Hill, NJ).

### Flow Cytometry

Hydrogels were divided into four regions, and identical regions from two separate hydrogels were combined into a single sample.^[14a]^ Hydrogels were degraded in a solution of 500 μL of PBS + 25% FBS and 100 Units Collagenase Type IV (#LS004186, Worthington Biochemical, Lakewood, NJ) for 30 minutes at 37 °C. The reaction was quenched in excess PBS and samples were centrifuged at 300 rcf x 10 minutes. The collected cells were resuspended in flow staining buffer (PBS + 5% FBS) and stained with surface marker antibodies for CD34, CD135, Lin, Sca1, and c-kit. Following surface staining, cells were prepped for intranuclear staining by fixation and permeabilization with the commercially available Foxp3/Transcription Factor Staining Buffer Set (#00-5523-00, Thermo Fisher Scientific, Waltham, MA), and subsequently stained for Ki-67 and DNA (DAPI; 1mg/mL, 10:300, #D21490, Thermo Fisher Scientific, Waltham, MA). Stained cells were resuspended in PBS + 5% FBS and analyzed via Fluorescence-Assisted Cytometry (FACs), using a BD LSR Fortessa (BD Biosciences, San Jose, CA). Fluorescent minus one (FMO) controls from lysed whole bone marrow were prepared fresh for each experiment for gating. DAPI was used to discriminate cells from debris, and analysis was performed with a 5,000 DAPI parameter thresholding. Cells were classified as Long-Term repopulating HSCs (LT-HSCs: CD34^−^ CD135^−^ Lin^−^ Sca1^+^ c-kit^+^);^[43]^ Short-Term repopulating HSCs (ST-HSCs: CD34^+^ CD135^−^ LSK);^[43]^ HSCs (CD34^+/-^ CD135^−^ LSK); or Multipotent Progenitors (MPPs: CD34^+^ CD135^+^ LSK).^[43c, 44]^ Cell cycle was classified as G0 (Ki-67^−^, DAPI^≤2N^); G1 (Ki-67^+^, DAPI^≤2N^); SGM (DAPI^>2N^). All antibodies were supplied by eBioscience (San Diego, CA), and are as follows: PE-conjugated Ki-67 (0.3:100, #12-5698-82), eFluoro660-conjugated CD34 (5:100, #50-0341-82), PE-CY5-conjugated CD135 (5:100, #15-1351-82), APC-efluor780-conjugated c-kit (1:160, #47-1172-81), PE-CY7-conjugated Sca-1 (0.3:100, #25-5981-81), and Lin: FITC-conjugated CD5, B220, CD8a, CD11b (1:100, #11-0051-82, #11-0452-82, #11-0081-82, #11-0112-82), Gr-1 (1:400, #11-5931-82), and Ter-119 (1:200, #11-5921-82).

### Immunofluorescent Staining

Hydrogels were fixed using formalin, permeabilized using 0.5% Tween 20, and blocked using 2% abdil or 5% donkey serum with 0.1% Tween 20. Mouse anti-human CD31 (1:200, #M0823, Agilent, Santa Clara, CA) or sheep anti-human CD31 (1:100, #AF806, R&D Systems, Minneapolis, MN) was used to stain for vascular networks, while goat anti-mouse CD45 (1:50, #20103-1-AP, Thermo Fisher Scientific, Waltham, MA) was used to identify hematopoietic cells. Chicken anti-mouse Alexa Fluor 488 (1:500; #A21200, Thermo Fisher Scientific, Waltham, MA), donkey anti-goat Alexa Fluor 555 (1:500; #A21432, Thermo Fisher Scientific, Waltham, MA), donkey anti-sheep NL637 (1:500; #NL011, R&D Systems, Minneapolis, MN) were used as secondary antibodies, and Hoescht 33342 (1:2000, 10 mg/ml, #H3570, Thermo Fisher Scientific, Waltham, MA) was used as a nuclear stain. Hydrogels were incubated with antibodies at 4 °C overnight and were washed with PBS + 0.1% Tween 20 between antibody incubation steps.

### Image Acquisition and Analysis

Fluorescent bead gradients were visualized using an AxioZoom v15 microscope (Zeiss, White Plains, NY). ImageJ (NIH, Bethesda, MD) was used to quantify mean fluorescent intensity across a hydrogel. Five equivalent regions of interest were drawn across the hydrogel, and mean fluorescent intensity was obtained for each region. Fluorescent intensity was normalized to the region with highest intensity after background subtraction.

To visualize vascular network formation as a function of cell or matrix density, hydrogels were divided into four regions. Each region was imaged for CD31+ microvasculature using a DMi8 Yokogawa W1 spinning disk confocal microscope with a Hamamatsu EM-CCD digital camera (Leica Microsystems, Buffalo Grove, IL). Z-stacks with a thickness of 200 μm and a step size of 5 μm were obtained for at least one area of interest per hydrogel region. Images containing vascular networks were skeletonized using the TubeAnalyst macro in ImageJ, and skeletons were quantified using a Matlab code described by Crosby et al. to extract metrics such as total network length, average branch length, number of branches, and number of junctions.^[45]^ All metrics except for average branch length were normalized to unit volume. If more than one area of interest was obtained for a hydrogel region, metrics were averaged across the areas of interest to obtain average values for the hydrogel region.

To visualize vascular network and hematopoietic cell association, hydrogels were divided into four regions. Each region was imaged using a Zeiss LSM 710 NLO (Zeiss, White Plains, NY). Z-stacks with a step size of 2.5 μm were obtained for at least one region of interest per hydrogel region. Maximum projection intensities were constructed from the z-stacks using Zeiss ZEN Black software. Representative images of each region were thresholded to maximize signal to noise.

### Western Blot

Individual hydrogels containing different cell densities were incubated on ice in RIPA buffer containing protease and phosphatase inhibitors (1:100, Sigma Aldrich, St. Louis, MO) for thirty minutes to lyse cells. Protein amounts were quantified using a BCA assay (Thermo Fisher Scientific, Waltham, MA). SDS-PAGE was performed under reducing conditions using a 10% Mini-PROTEAN TGX precast polyacrylamide gel (Bio-Rad, Hercules, CA) with 5 μg protein per sample. Subsequently, proteins were transferred onto nitrocellulose membranes (GE Healthcare Bio-Sciences, Pittsburgh, PA). Milk or bovine serum albumin in TBST was used for blocking and dilution of antibodies. Primary antibodies included total p42/44 ERK1/2 (1:1000, #4695), phosphorylated p42/44 ERK1/2 (1:2000, #4370), total Akt (1:1000, #4691), phosporylated Akt(S473) (1:2000, #4060), and β-actin (1:1000, #4970), while horseradish peroxidase-conjugated anti-rabbit (1:2000, #7074) was used as the secondary antibody (Cell Signaling Technology, Danvers, MA). Membranes were stripped using OneMinute Western Blot Stripping Buffer (GM Biosciences, Frederick, MD) for one minute. The SuperSignal West Pico Chemiluminescent Substrate kit (Thermo Fisher Scientific, Waltham, MA) was used for detection by an ImageQuant LAS 4000 (GE Healthcare Bio-Sciences, Pittsburgh, PA). Band intensities were analyzed using ImageJ by subtracting the average band intensity from the average background intensity, and phosphorylated protein amounts were normalized to the total protein amounts for each sample. Five hydrogel samples were assessed per condition.

### Cytokine Array

Conditioned media from individual hydrogels with varying vascular density was analyzed using the Proteome Profiler Human XL Cytokine Array Kit (R&D Systems, Minneapolis, MN). 0.5 mL of media was used per array, and three media samples were assessed per condition. Cytokine spot intensities were obtained using an ImageQuant LAS 4010 (GE Healthcare Bio-Sciences, Pittsburgh, PA). ImageJ was used to quantify the intensities of the cytokine spots, which were normalized to the intensities of the positive reference spots on each array.

### Statistics

Statistics were performed using OriginPro (OriginLab, Northampton, MA). Normality of data was determined using the Shapiro-Wilk test, and equality of variance was determined using Levene’s test. For normal data, comparisons between multiple groups were performed using a one-way ANOVA with Tukey’s post-hoc when assumptions were met. In the case where data was not normal or groups had unequal variance, comparisons between multiple groups were performed using a Kruskal-Wallis test with Dunn’s post-hoc. Significance was determined as p < 0.05. Data is reported as mean ± S.D., and sample size is reported in the relevant materials and methods or in the figure caption.

## Acknowledgements

The authors would like to acknowledge Dr. Barbara Pilas and Basia Balhan from the Flow Cytometry Unit at the Roy J. Carver Biotechnology Center (UIUC) for their assistance and technical expertise. Additionally, the authors would like to thank the Carl R. Woese Institute for Genomic Biology for providing core microscopy and imaging facilities that were instrumental for the data generated in this study. Funding sources include the NIH (R01 CA197488, F31 DK117514, T32 EB019944, R01 DK099528, R21 EB01848) and NSF (GRFP DGE-1144245). The content is solely the responsibility of the authors and does not necessarily represent the official views of the NIH or NSF. The authors are also grateful for additional funding provided by the Department of Chemical & Biomolecular Engineering, the Carl R. Woese Institute for Genomic Biology, and the Illinois Scholars Undergraduate Research Program at the University of Illinois at Urbana-Champaign.

## Supporting Information

**Supplementary Figure 1.**
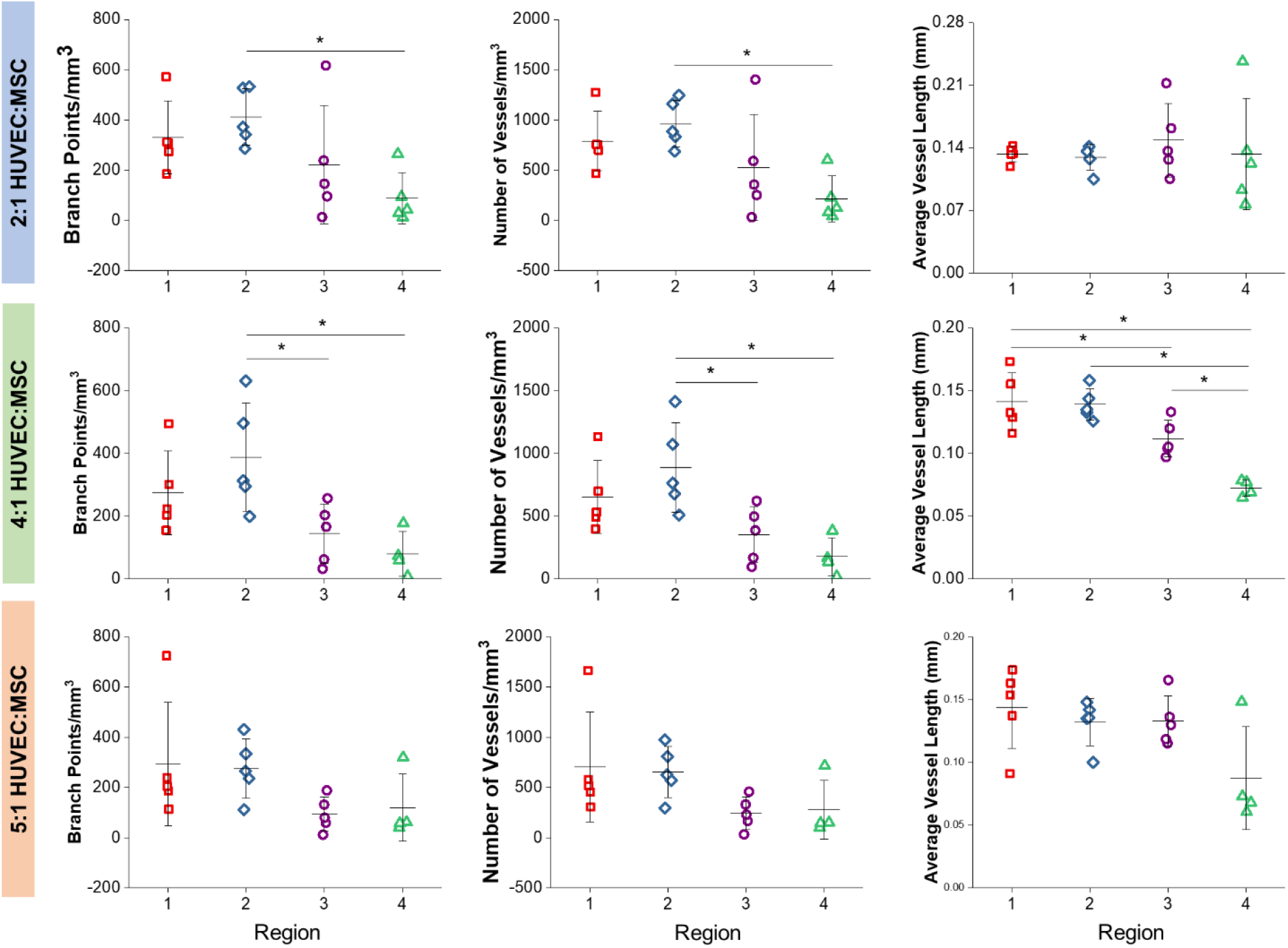
Additional metrics for vascular complexity reveals that gradient resolution is best achieved with 2:1 and 4:1 HUVEC:MSC ratios. *p<0.05, N = 5 devices.

**Supplementary Figure 2.**
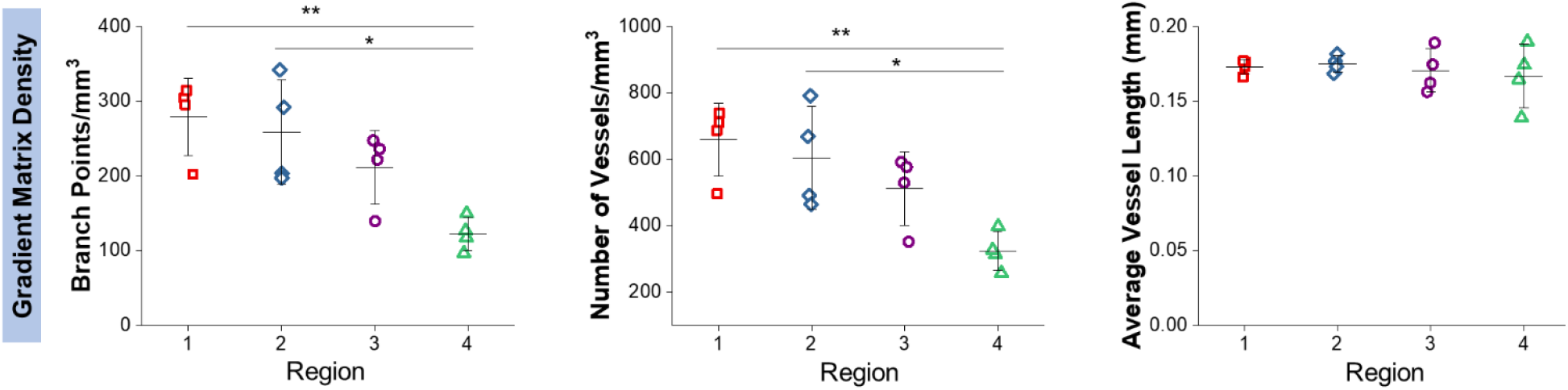
Additional metrics for vascular complexity reveals that a gradient in matrix density generates regional differences in vascular architecture. *p<0.05, **p<0.01, N = 4 devices.

**Supplementary Figure 3.**
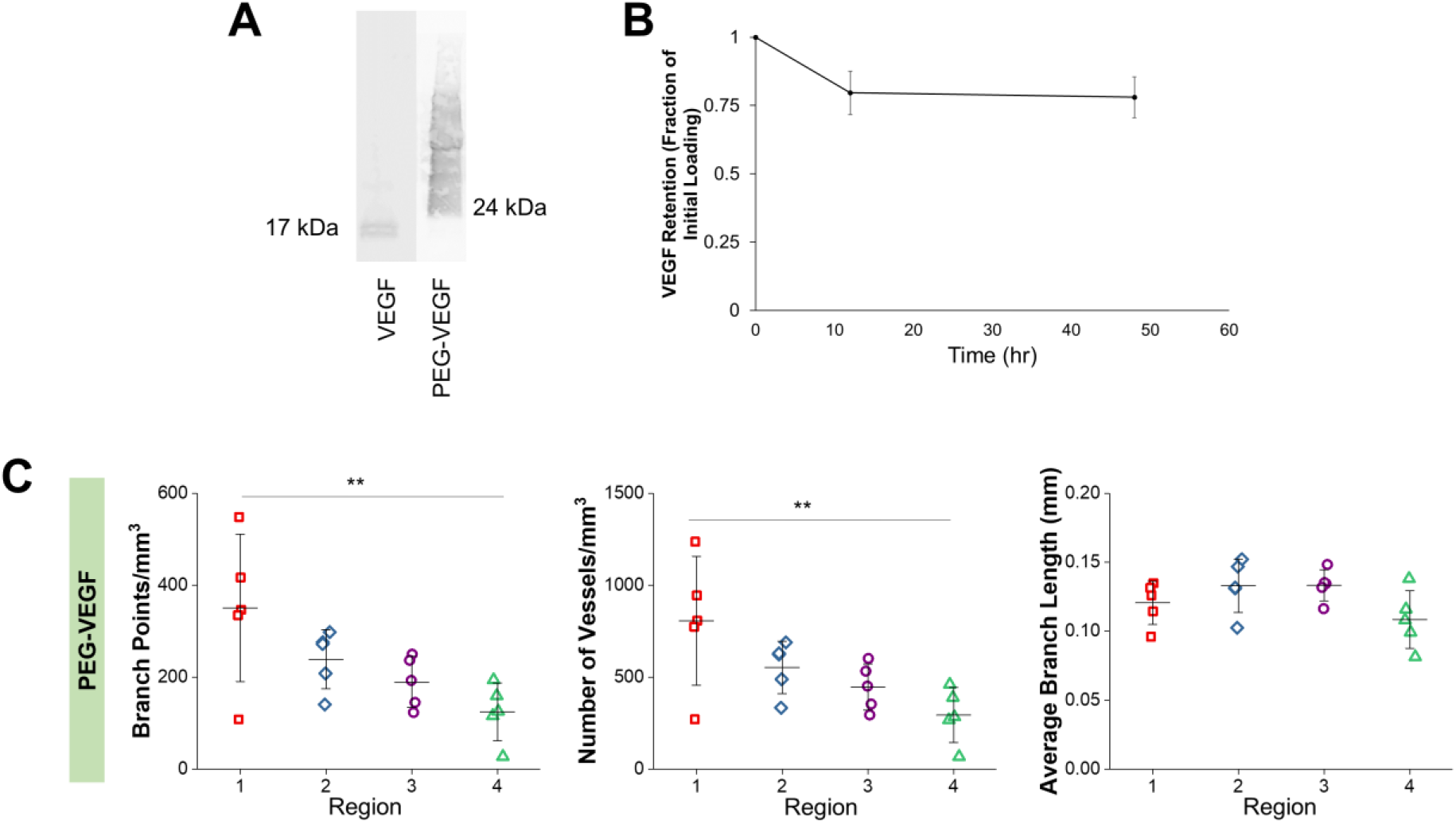
**(A)** Western blot membrane demonstrating the successful conjugation of VEGF to acrylate-PEG-succinimidyl ester. **(B)** Retention of covalently-bound VEGF in GelMA over 48 hours. **(C)** Additional metrics for vascular complexity reveals that a gradient in PEG-VEGF generates regional differences in vascular architecture. **p<0.01, N = 5 devices.

**Supplementary Figure 4.**
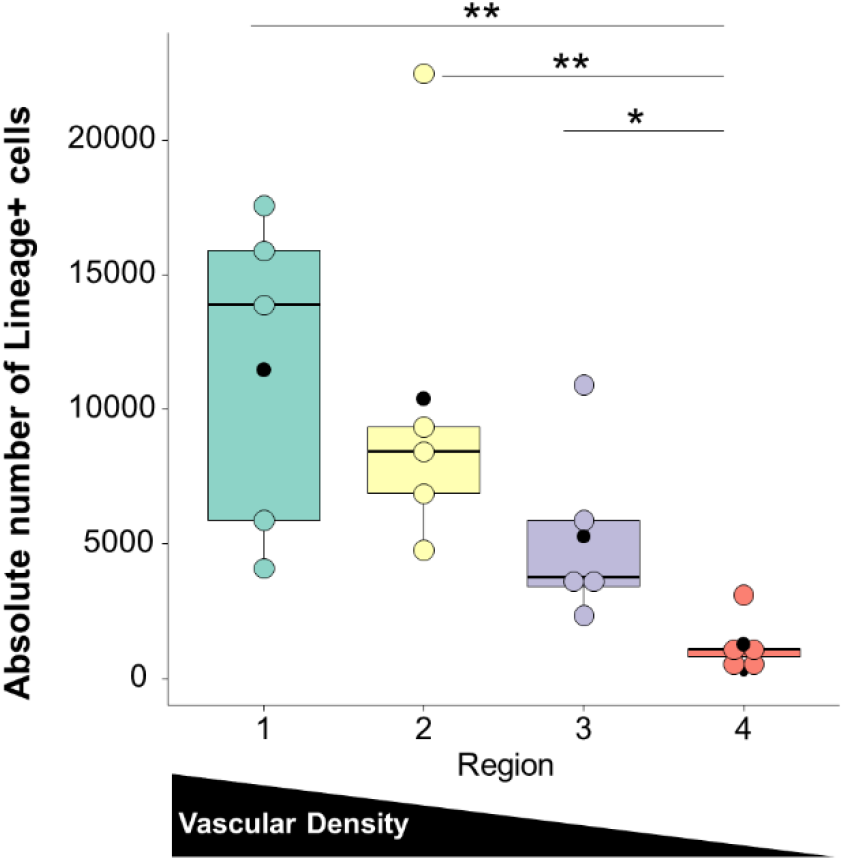
The number of lineage-positive hematopoietic cells significantly increased with increasing vascular density. The graph depicts the absolute number of Lin+ hematopoietic cells in each region. *p<0.05, **p<0.001 N = 5 devices.

**Supplementary Figure 5.**
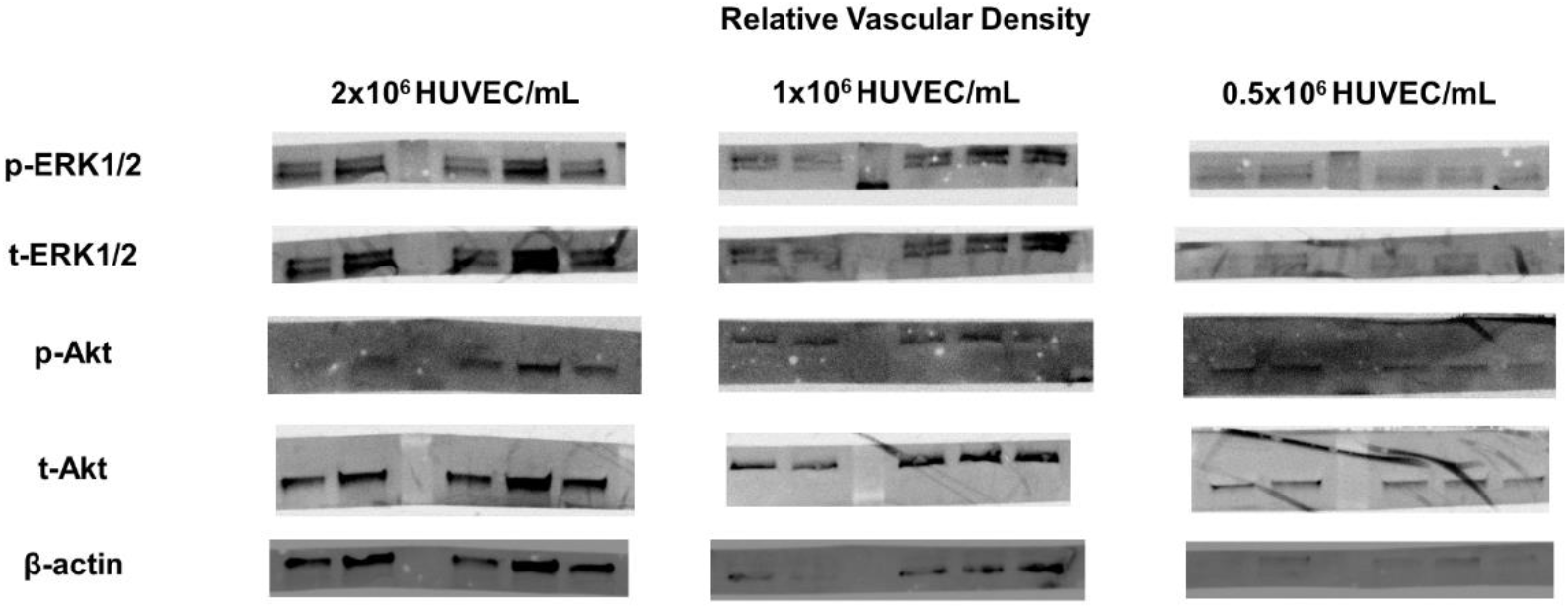
Images of Western blot membranes.

